# Lineage does not regulate the connectivity of projection neurons in the mouse olfactory bulb

**DOI:** 10.1101/584805

**Authors:** Luis Sánchez-Guardado, Carlos Lois

## Abstract

Lineage regulates the synaptic connections between neurons in some regions of the invertebrate nervous system. In mammals recent experiments suggest that cell lineage determines the connectivity of pyramidal neurons in the neocortex, but the functional relevance of this phenomenon and whether it occurs in other neuronal types remains controversial. We investigated whether lineage plays a role in the connectivity of mitral and tufted cells, the projection neurons in the mouse olfactory bulb. We used transgenic mice to label neuronal progenitors sparsely and observed that clonally related neurons receive synaptic input from olfactory sensory neurons expressing different olfactory receptors. These results indicate that lineage does not determine the connectivity between olfactory sensory neurons and olfactory bulb projection neurons.

## Introduction

The relationship between cell lineage and neuronal connectivity in the brain is not well understood. Lineage regulates the synaptic connections between neurons in some regions of the invertebrate nervous system. For example, in the *Drosophila* olfactory system, projection neurons are specified by cell lineage to receive synaptic input from the axons of specific types of olfactory sensory neurons (OSNs) (Jefferis et al., 2001; Li et al., 2018). In mammals it has been reported that clonally related pyramidal neurons are preferentially connected to each other in the neocortex (Yu et al., 2009; 2012; He et al., 2015). Furthermore, it has been proposed that sister neurons in the visual cortex have a strong correlation to the stimuli to which they respond (Li et al., 2012), while other works suggest that this correlation is much weaker (Ohtsuki et al., 2012). To further investigate the role played by lineage in the assembly of brain circuits we focused on the mammalian olfactory bulb, a brain region with an anatomical organization particularly advantageous to study this question.

The mammalian olfactory system can be divided into three regions: olfactory epithelium, olfactory bulb (OB) and olfactory cortex. The olfactory epithelium harbors the OSNs. Each OSN expresses just one of more than one thousand odorant receptors (Buck and Axel, 1991; Chess et al., 1994). OSN axons expressing the same odorant receptor converge into one or two discrete neuropil structures in each OB called glomeruli, forming a stereotypic map on the OB surface (Ressler et al., 1994; Vassar et al., 1994; Mombaerts et al., 1996; Wang et al., 1998). The projection neurons in the OB are called mitral and tufted cells (M/T cells). In mammals the majority (>90%) of M/T cells have a single apical dendrite that branches into a single glomerulus (Mori, 1987; Shepherd and Shepherd, 1990; Malun and Brunjes, 1996) where they receive sensory input from OSNs expressing a particular odor receptor (Figure 1A) (Ressler et al., 1994; Vassar et al., 1994). Thus, the anatomical organization of the glomerulus in the OB is an ideal system to investigate the possible relationship between lineage and connectivity because the apical dendrite of the M/T cells provides a direct readout of their synaptic input. To address this question we sparsely labeled M/T cells progenitors and investigated the sensory input that their progeny receives from OSNs. Our results show that sister M/T cells receive synaptic input from different glomeruli, indicating that lineage does not determine the neuronal connectivity of the OB projections neurons, and suggest that the assembly of the OB mostly depends on non-genetic mechanisms.

**Figure 1.**
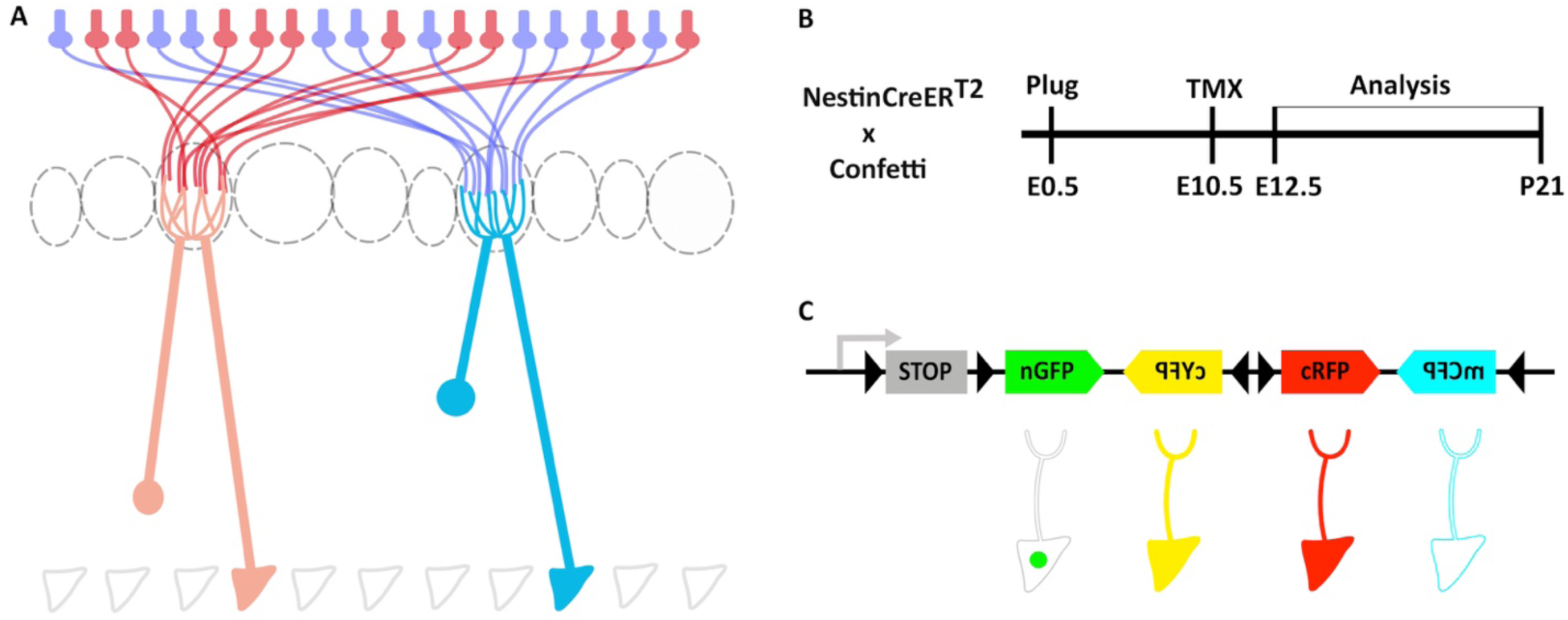
Clonal analysis of projection neurons using *Nestin-CreER*^*T2*^::*Confetti* mice to sparsely label neuronal progenitors. (A) Schematic representation of the olfactory bulb (OB). Axons from olfactory sensory neurons (OSNs) expressing the same receptor project to a single glomerulus, forming synaptic contacts with the apical dendrites of mitral and tufted cells. (**B**) Experimental design to label neuronal progenitors with tamoxifen (TMX) at embryonic day 10 (E10.5) and their posterior analysis at E12.5 and P21. (**C**) The *Confetti* cassette encodes 4 different fluorescent proteins (nuclear GFP (nGFP), membrane CFP (mCFP), and cytoplasmic YFP (cYFP) and RFP (cRFP)). Upon Cre recombination, the STOP sequence is excised and randomly generates four possible outcomes.

## Results and discussion

### Labeling of progenitors of OB projection neurons

The projection neurons in the OB are called mitral and tufted cells (M/T cells). M/T cells originate from progenitors located in the OB primordium, which is derived from the anterior part of the dorsal telencephalon (Hinds, 1968a, 1968b). To investigate the lineage of M/T cells, we crossed two transgenic mouse, *Nestin-CreER*^*T2*^ (Kuo et al., 2006), which can be used to label neuronal progenitors in a sparse manner, with the *Confetti* line (Snippert et al., 2010), which can label individual cells with one out four possible fluorescent proteins upon Cre-mediated recombination (Figure 1B, C and Figure 1-figure supplement 1).

In order to optimize the conditions to label just a handful of progenitors, ideally a single progenitor per OB, we performed some preliminary experiments. First, we confirmed that our transgenic mice *Nestin-CreER*^*T2*^::*Confetti* did not label any neurons in the brain without tamoxifen (TMX) administration (n=3; data not shown). Second, we found that with an injection of 1 mg of TMX per 40 grams of body weight into a 10-day pregnant female (E10.5) we observed a handful of pyramidal neuron clones in the neocortex, and around 20 M/T cells labeled in the OB when the brains were examined at postnatal day 21 (P21) (Figure 1B and and Figure 1-figure supplement 1). Third, we confirmed that this TMX concentration labeled a few progenitors per brain when animals were analyzed two days after TMX administration (E12.5). With these conditions, we observed between none to a single progenitor labeled per fluorescent protein in the OB (n=6) (Figure 2). Although we observed a very low number of progenitors labeled, we cannot determine whether a group of cells labeled at P21 with the same fluorescent protein in the OB originated from a single progenitor, or from two independent progenitors. However, here we will work under the assumption that any group of M/T cells labeled with the same fluorescent protein in the OB are part of a single clone.

**Figure 2.**
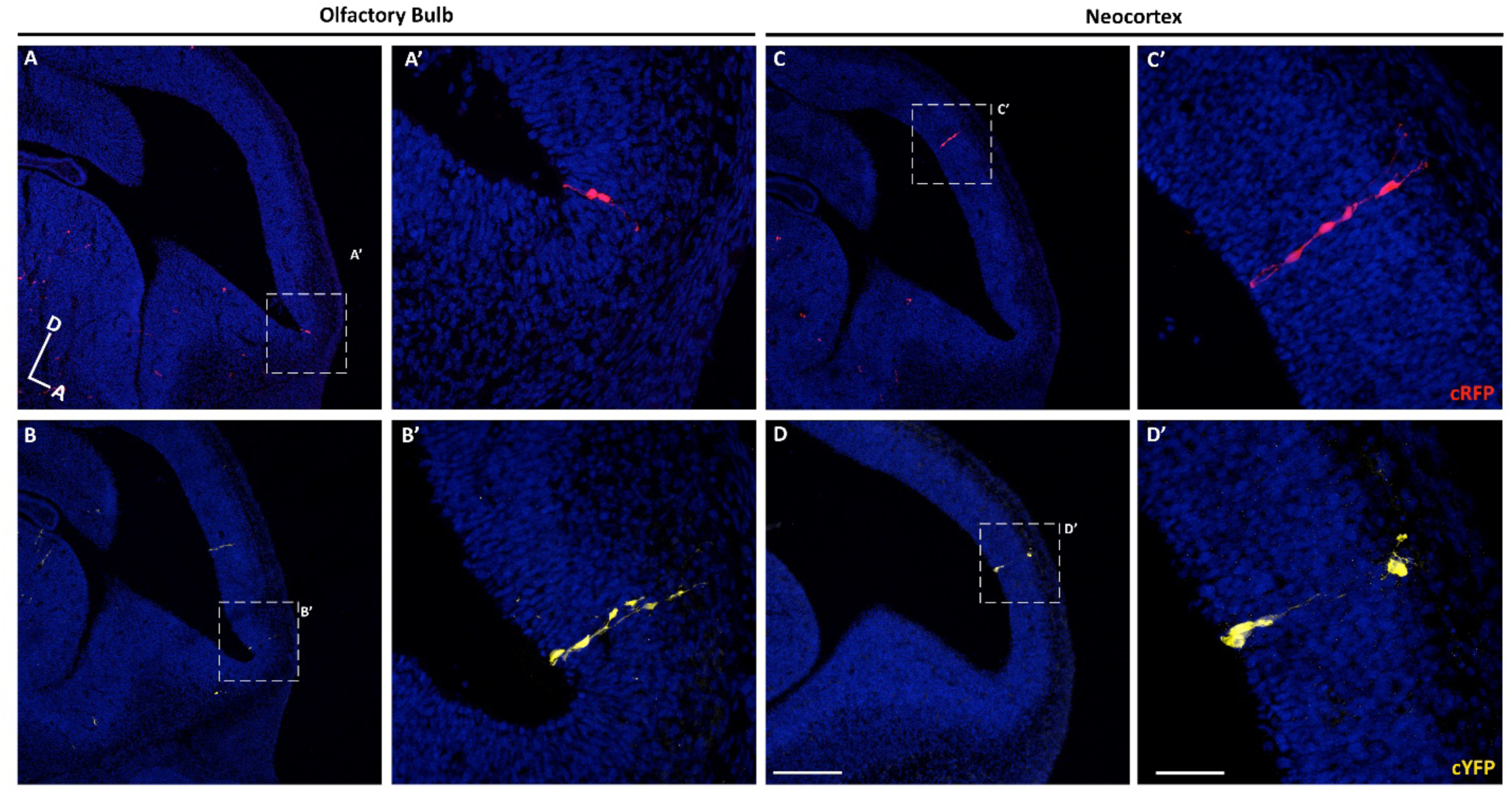
Sparse labeling of progenitor cells in the embryonic mouse brain. (**A-D**) Sagittal sections through the brain of an E12.5 mouse treated with TMX at E10.5. (**A-B**) Confocal images of individual clones labeled in the OB expressing cRFP (**A-A’**) and cYFP (**B-B’**). (**A’**-**B’)** High magnification images of the clones showed in A and B. (**C-D**) Single clones labeled in the neocortex expressing cRFP (**C-C’**) and cYFP (**D-D’**). (**C’-D’**) High magnification images of the clones showed in **C-D**. Cell nuclei are labeled with DAPI (blue). Scale bar in D is 200 µm and applies to A-D, scale bar in D’ is 50 µm applies to A’-D’. Orientation of brains: D, dorsal; A, anterior.

To study the lineage of the M/T cells we induced Cre activity at E10.5, the peak time for mitral cell generation (Hinds, 1968a, 1968b; Blanchart et al., 2006; Kim et al., 2011; Imamura et al., 2011). Brains were analyzed at P21, once M/T completed the refinement of their dendrites and they have a mature morphology with a single apical dendrite projecting into a single glomerulus (Figure 1A) (Malun and Brunjes, 1996; Lin et al., 2000; Matsutani and Yamamoto, 2000; Blanchart et al., 2006). *Confetti* mice can produce four different fluorescent proteins with distinct subcellular locations (cytosolic (cRFP and cYFP), membrane (mCFP), and nuclear (nGFP)) (Figure 1C, Figure 1-figure supplement 1 and Figure 2-figure supplement 1) (Snippert et al., 2010). Consistent with previous works, we observed that the majority of clones in the OB were labeled by RFP (n=9), whereas YFP (n=4) and CFP (n=1) clones appeared less frequently (Reeves et al., 2018). However, we did not analyze any of the nGFP+ cells for two reasons. First, the most reliable way to unambiguously identify M/T cells is by their distinctive morphology. However, if a cell is only labeled in the nucleus (as in nGFP+ cells), we cannot tell apart M/T cells from other OB cell types (e.g., short axon cells, granule cells, juxtaperiglomerular). Second, to identify the connectivity between M/T cells and glomeruli, it is necessary to follow the projection of their apical dendrites (Figure 1-figure supplement 1), and we cannot observe any dendrites in the nGFP+ cells.

In total, we analyzed 29 OBs, and 13 of them did not have any labeled cells. Out of the 16 OBs with labeled cells, 14 OBs had both M and T cells, and 2 OBs had only M cells labeled (with 3 and 4 M cells labeled per OB). We do not know the reason why these two OBs showed only M cells, and several reasons may account for this observation, including progenitors committed to produced only M cells, or labelling of intermediate progenitor that underwent few cell divisions. We did not find any OB with only T cells.

### Size of clones and distribution of neurons in the OB and neocortex

We measured the putative clone size in the OB and compared them with neocortex clones. We found that putative clones in the OB contained 22.14 ± 6.61 M/T cells (average ± standard deviation, n= 310), while neocortex clones contained 92.67 ± 23.18 pyramidal neurons (n=556), consistent with previous results (Franco et al., 2012; Gao et al., 2014) (Figure 3A). These observations suggest that the clone size in the neocortex is four times larger than a clone in the OB, consistent with the reported different modes of neurogenesis in each of these two brain regions (Cárdenas et al., 2018).

**Figure 3.**
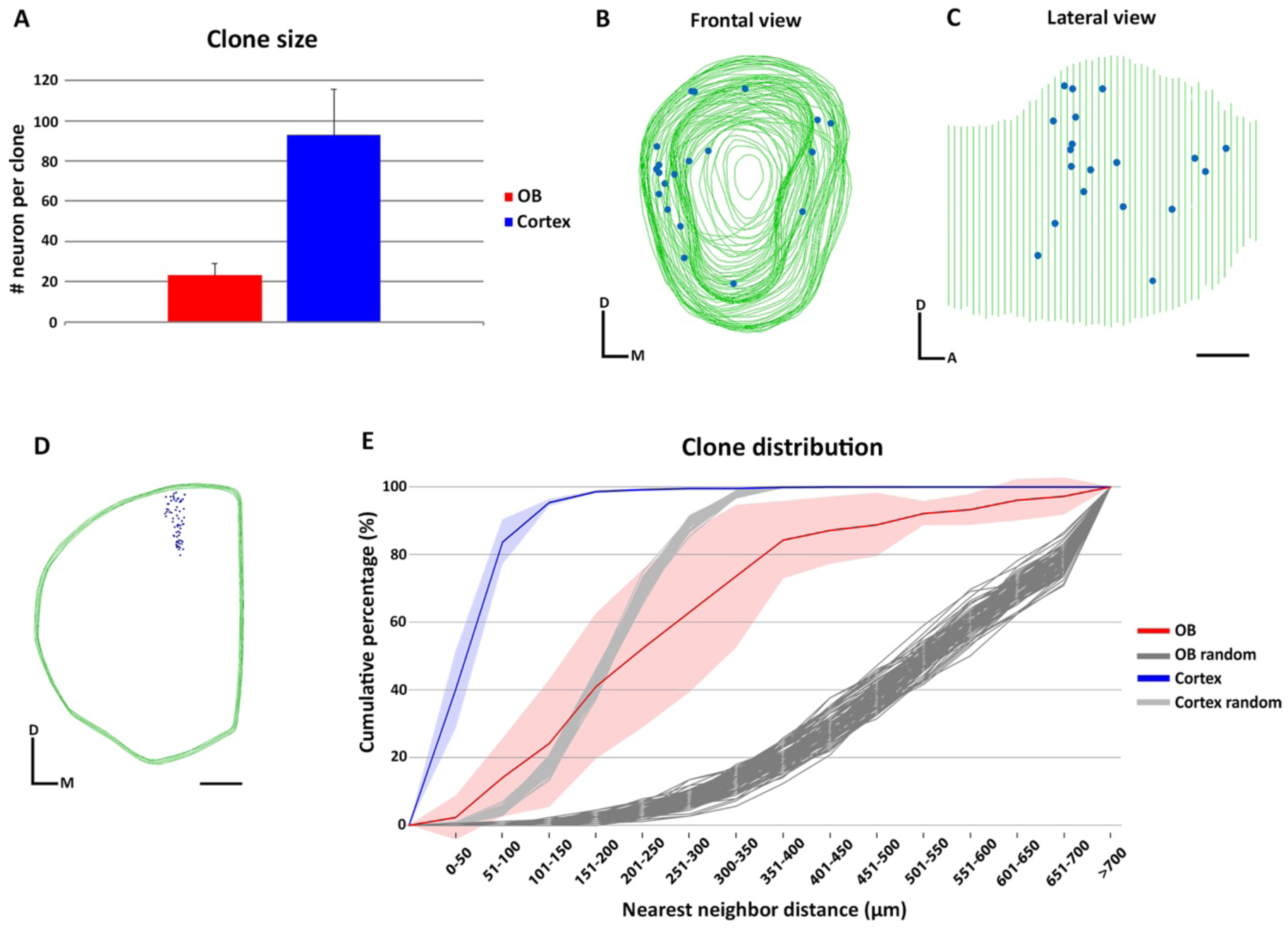
Clone size and distribution of cells labeled in the olfactory bulb and neocortex. (A) Clone size quantification in the OB and neocortex. Data are shown as average ± standard deviation. (**B-D**) 3D reconstruction of a *NestinCreER*^*T2*^::*Confetti* P21 mice OB (**B-C**) and neocortex (**D**) treated with TMX at E10.5. Green lines indicate the contours of the brain and blue dots represent the cell bodies of labeled neurons. (**B**) Frontal and (**C**) lateral views of the 3D reconstruction of one OB. (**D**) Frontal view of the neocortex 3D reconstruction. (**E**) Cumulative percentage of the NNDs of sister neurons labeled in the OB (red) and neocortex (blue). Data are shown as average ± standard deviation (OB, n=178 neurons in 9 clones; neocortex, n=556 neurons in 6 clones). Dark and light gray lines represent 100 datasets of random simulations of OB and neocortex NND, respectively. Scale bar in C is 0.5 mm and applies to B-C. Scale bar in D is 1 mm. Orientation of diagrams in B-D: D, dorsal; A, anterior; M, medial.

We analyzed the distribution of cell bodies of 9 OB clones (n=178 neurons) and 6 neocortex clones (n=556 neurons) by performing 3D reconstructions using the Neurolucida software (Figure 3B-D and Figure 3-figure supplement 1). The 3D reconstructions revealed that sister M/T cells were distributed in a broader area than the tight columns of sister pyramidal neurons. To analyze the distribution of cells from each clone, we calculated the nearest neighbor distance (NND) based in our 3D reconstructions using Neurolucida Explorer (Figure 3E and Figure 3-figure supplement 2). We found that sister M/T cells were more separated from each other (283.96 µm ± 70.28; average ± standard deviation) than sister pyramidal neurons (65.45 µm ± 19.4) (Figure 3E). The distribution of sister M/T cells that we observed is consistent with the tangential migration of immature M/T cells reported in the embryonic OB (Blanchart et al., 2006; Imamura et al., 2011).

To investigate whether the distribution of sister M/T cells observed was random, we compared the NNDs of the labeled M/T cells observed (n=178) with a simulated random dataset. The same strategy was followed for neocortex clones. We found that the NNDs between clonally related neurons were shorter than the simulated random datasets both for the OB and neocortex (Figure 3E). Similar results were reported for pyramidal neurons in the neocortex (Gao et al., 2014). This indicates that although sister M/T cells are not clustered as pyramidal neurons, their distribution in the OB is not random. Interestingly, a previous work have observed that the tangential migration of immature M/T cells in the embryonic OB may be regulated by gradients of secreted molecules, limiting their distribution to specific regions within the OB (Inokuchi et al., 2017).

### Connectivity of sister M/T cells

It has been proposed that the anatomical organization of the OB may be analogous to the neocortex columnar organization. In the neocortex it is thought that the pyramidal neurons forming part of a column perform a similar task (Mountcastle, 1997). Similarly, M/T cells receiving synaptic input from the same glomerulus may also perform a similar task (Kauer and Cinelli, 1993; Mori et al., 1999; Bozza et al., 2002). Our results indicate that sister M/T cells are widely distributed throughout the OB (Figure 3). Based on this observation, it seems unlikely that sister M/T cells would have apical dendrites projecting into the same glomerulus. Although improbable, this could still be possible because the soma of M/T cells innervating the same glomerulus may be separated from each out up to 450 µm (for M cells) and 350 µm (for T cells) (Liu et al., 2016). To investigate whether sister M/T cells receive synaptic input from the same glomerulus, we tracked their apical dendrites (Figure 4). Among all the labeled M/T cells that we detected (n=310, from 14 putative M/T clones) we never observed two neurons innervating the same glomerulus, even when their cell bodies were near each other (Figure 4B-E). Nevertheless, it is still possible that, although we did not observe them, there may exist clones of M/T cells genetically pre-determined to project to the same glomerulus. This scenario could be expected for putative glomeruli responsive to relevant odors for survival, such as those responsive to predators or poisons, which require an innate and hardwired response of avoidance (Sosulski et al., 2011). Future experiments analyzing a much larger number of clones than those detected here may reveal the existence of these putative “hardwired” M/T clones.

**Figure 4.**
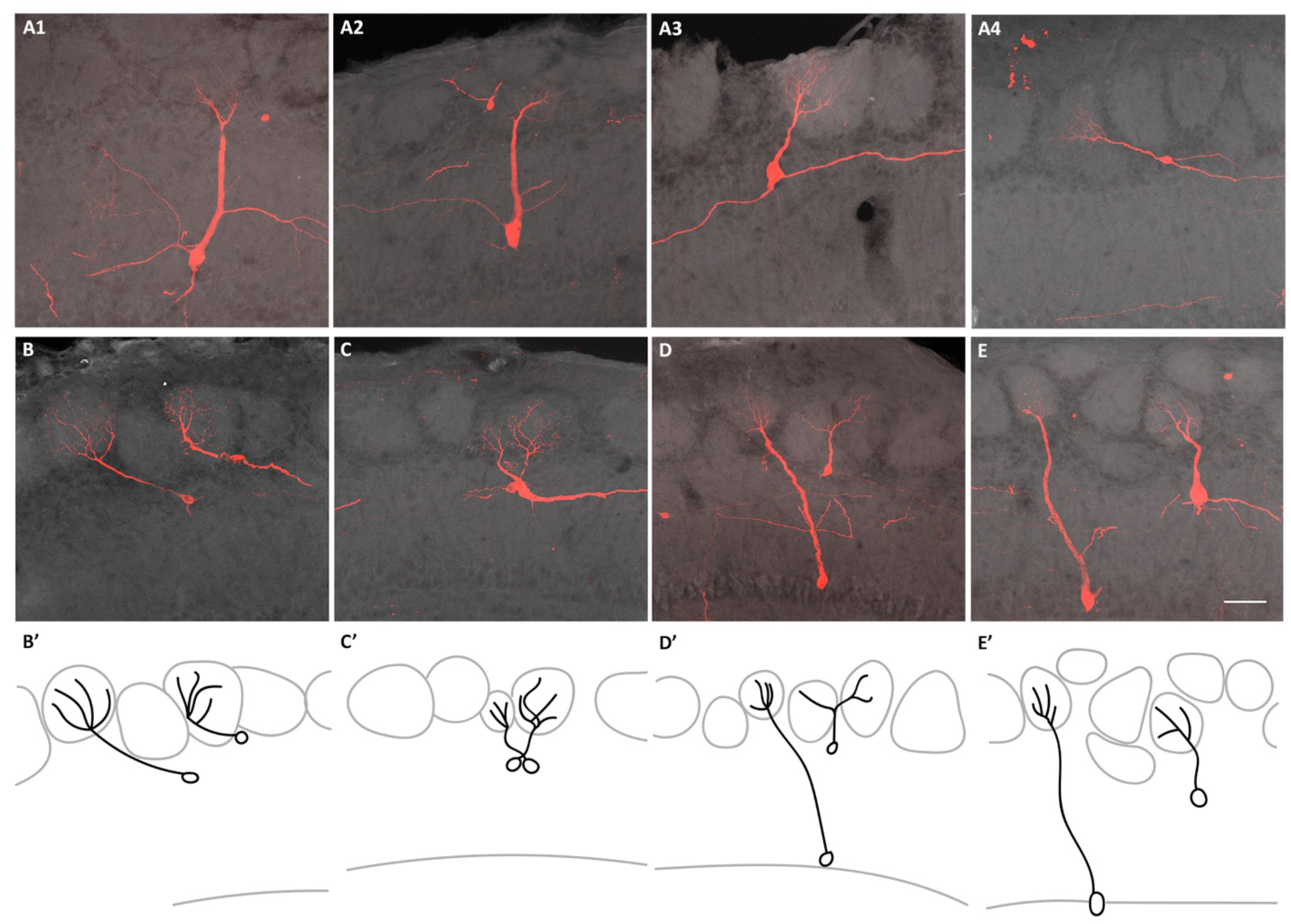
Connectivity clonally related M/T cells. (**A**) Confocal images of four sister M/T cells belonging to a putative individual clone in the OB (**B-E**) Confocal images of sister M/T cells from four clones, in four different OBs, with their somata close to each other and their apical dendrites innervating different glomeruli. (**B’-E’**) Schematic representation of the confocal images in B-E. Scale bar in E is 50 µm and applies to A-E.

In summary, our results indicate that lineage does not determinate the input connectivity of the projection neurons in the mammalian OB. This is in contrast to what has been described for projection neurons in the *Drosophila* antennal lobe (Jefferis et al., 2001) and suggested for pyramidal neurons in the rodent visual cortex (Li et al., 2012). Our results suggest that the sensory input received by M/T cells is regulated by non-genetic factors, consistent with the observations from recent works. For example, it has been shown that sensory odor experience starting *in utero* recruits the apical dendrites of M/T cells to the activated glomeruli (Liu et al., 2016).

Is there any biological advantage to the dispersion of sister projection neurons in the OB? Interestingly, it has been proposed that the M/T cells receiving input from the same glomerulus exhibit a wide diversity in their biophysical properties, and this diversity may be important for neural coding (Padmanabhan and Urban, 2010). In addition, neurons in the piriform cortex receive synaptic input from M/T cells innervating different glomeruli (Miyamichi et al., 2011), whereas M/T cells connected to the same glomerulus project their axons into many different areas of the olfactory cortex (Ghosh et al., 2011; Sosulski et al., 2011). However, the connectivity between M/T cells and the amygdala appears to be more stereotypical than between the M/T cells and other targets in the olfactory cortex (anterior olfactory nucleus, piriform cortex, tenia tecta, olfactory tubercle, cortical amygdala and entorhinal cortex) (Haberly, 2001; Sosulski et al., 2011). Based on these observations, one can speculate that the connectivity between the OB and its targets in the olfactory cortex may occur by two different mechanisms. Genetic factors, including lineage, may contribute to the connectivity between M/T cells and the amygdala, as this brain area is involved in innate behavior responses that may require hardwired connections (Sosulski et al., 2011). In contrast, the connectivity between M/T cells and areas of the olfactory cortex involved in the perception of odors that do not elicit innate behaviors are more plastic and may be regulated by non-genetic mechanisms, such as activity-dependent wiring, among others (Caron et al., 2013; Schaffer et al., 2018).

Our results indicating that lineage does not determine the synaptic input of M/T cells raise further questions about the assembly of the olfactory circuits, including which are the mechanisms that regulate the connectivity between M/T cells and OSNs, the role that experience may play sculpting the odor representations in the piriform cortex, and whether lineage regulates the connections with the amygdala to trigger innate behaviors.

## Materials and Methods

### Animals

*Nestin-CreER*^*T2*^ and *Confetti* mice were obtained from Jackson Laboratory. The *Nestin-CreER*^*T2*^ mice can be used to induce the activity of Cre recombinase in neuronal progenitors by the administration of tamoxifen (TMX) into animals (Kuo et al., 2006). The *Confetti* mouse is a Cre-dependent reporter that produces four different fluorescent proteins (Snippert et al., 2010). We crossed the *Nestin-CreER*^*T2*^ mouse with the *Confetti* mice, and the resulting transgenic *Nestin-CreER*^*T2*^::*Confetti* mouse was used for the experiments. For the timed pregnancy, the plug date was designated as E0.5 and the day of birth as P0. In all experiments, mice were handled according to the protocols approved by the Caltech Institutional Animal Care and Use Committee (IACUC). Mice colonies were maintained at the animal facility of the California Institute of Technology (Caltech).

### Tamoxifen induction

Tamoxifen (TMX, Sigma T-5648) was dissolved in 37°C pre-warmed corn oil (Sigma C8267) at a concentration of 10 mg/ml. *NestinCreER*^*T2*^::*Confetti* embryos were induced at E10.5 (embryonic day 10.5) by a single intraperitoneal injection of 1 mg TMX into pregnant females (∼ 40 grams). Animals were euthanized at embryonic day 12 (E12.5) or postnatal day 21 (P21).

### Tissue processing, immunohistochemistry, and imaging

Mouse embryos (E12.5) were fixed by immersion in 4% paraformaldehyde (PFA) in phosphate-buffered saline (PBS, pH 7.4) at 4°C overnight. Postnatal mice (P21) were fixed by intracardiac perfusion with 4 % PFA in PBS. Brains were then extracted and incubated in 4% PFA at 4°C overnight. Next day, all samples were washed 3 times, 10 minutes each, with 0.1 M PBS, pH 7.4. Postnatal mice brains were embedded into 3 % agarose and cut in a vibratome into 60 µm thick sections. Sections were collected sequentially. Embryonic brains were cut with a cryostat into 20 µm thick sections as previously described (Sánchez-Guardado et al., 2009).

We amplified the signal from fluorescent proteins by performing immunohistochemistry with antibodies against RFP and GFP. Although anti-GFP antibody recognizes nGFP, cYFP and mCFP proteins, we were able to distinguish between them based on the different subcellular location of the proteins (nuclear, cytoplasmic and membrane). In the figures cells are shown with their original colors from the *Confetti* cassette, even though the signal from cYFP and mCFP proteins was amplified using the antibody against GFP (Figure 1-figure supplement 1, Figure 2, Figure 2-figure supplement 1). We did not include nGFP+ cells in our analyses because we cannot identify their morphology.

For immunocytochemistry, we incubated the sections during 30 minutes in blocking solution containing 1% bovine serum albumin in 0.1 M PBS-0.1% Triton X-100 (PBS-T). Sections were incubated overnight with the following antibodies diluted into blocking solution: chicken anti-GFP (1:1,000; AB3080; Millipore Bioscience Research Reagents), rabbit anti-RFP (1:1,000; LS-C60076; Lifespan). The next day sections were washed 3 times, 10 minutes each, in PBS-T. Later, sections were incubated during 90 minutes at room temperature with secondary antibodies (Alexa Fluor 488 goat anti-rabbit, Alexa Fluor 555 goat anti-chicken; Invitrogen) diluted 1:1,500 in blocking solution. Finally, the sections were counterstained with DAPI (D9542, Sigma), mounted sequentially on glass slides and mounted with Fluoromount (F4680, Fluoromount Aqueous Mounting Medium).

Z-stacks images were acquired using10x, 20x or 40x objectives on a confocal microscope (Zeiss LSM 800). Z-stacks were merged and analyzed using ImageJ and edited with Photoshop (Adobe) software.

### 3D reconstruction and data analysis

Each section was analyzed and traced in sequential order from rostral to caudal using Neurolucida and StereoInvestigator software (MBF Bioscience Inc., Williston, VT). The boundaries of the OB and neocortex were traced and used to line up each section with the previous one to form 3D reconstructions. Each labeled cell in the OB or neocortex was tagged with a blue dot.

The distribution of the nearest neighbor distance (NND) was calculated using Neurolucida Explorer software based on our 3D reconstruction. NND was calculated by identified the shortest straight path between labeled cells. The NND was represented as cumulative percentage (average ± standard deviation) of the clones analyzed in the OB (n=9) and neocortex (n=6). In addition, we generated a dataset of random simulations based on the same number of the M/T cells detected in our experiments (n=178). The distances were generated randomly with a normal distribution between the longest and shortest distances observed between M/T cells (closest and farthest sister M/T cells were separated by 21.51 µm and 974.82 µm, respectively), and repeated 100 times. We followed the same procedure for pyramidal neurons (n=556) in the neocortex (closest and farthest sister pyramidal neurons were separated by 13.54 µm and 415.2 µm, respectively).

## Acknowledgements

We are grateful to Walter G. Gonzalez and Antuca Callejas for comments on the manuscript.

---

**Figure 1-figure supplement 1.**
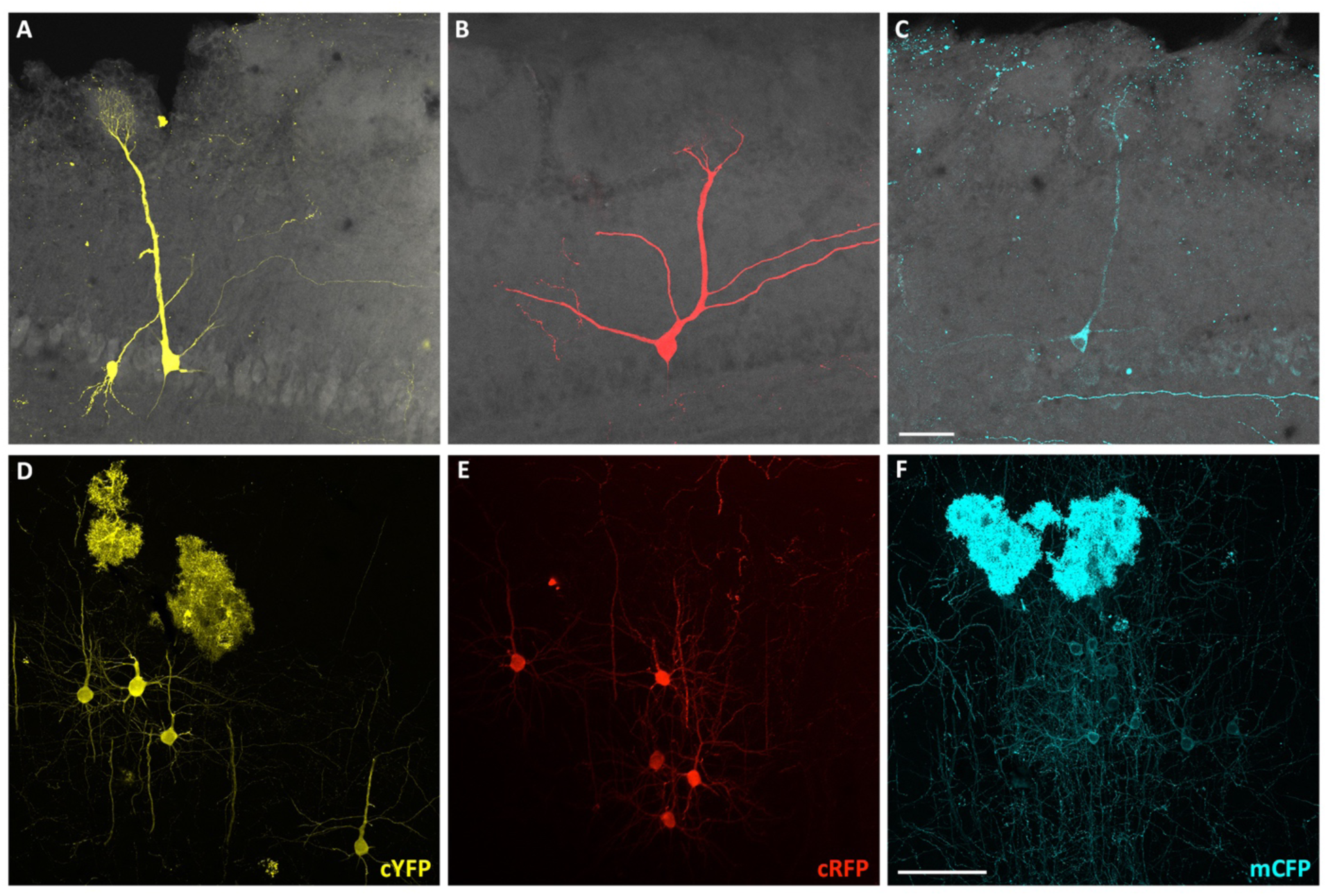
Pyramidal and M/T neurons labeled with different fluorescent proteins. (**A-C**) Confocal images of three M/T cells and (**D-F**) three pyramidal neuron clones labeled with different fluorescent proteins in OB and neocortex coronal sections of P21 mice treated with TMX at E10.5. (**A, D**) Cytoplasmic YFP (cYFP); (**B, E**) cytoplasmic RFP (cRFP) and (**C, F**) membrane CFP (mCFP). Scale bar in C is 50 µm. Scale bar in F is 100 µm.

**Figure 2-figure supplement 1.**
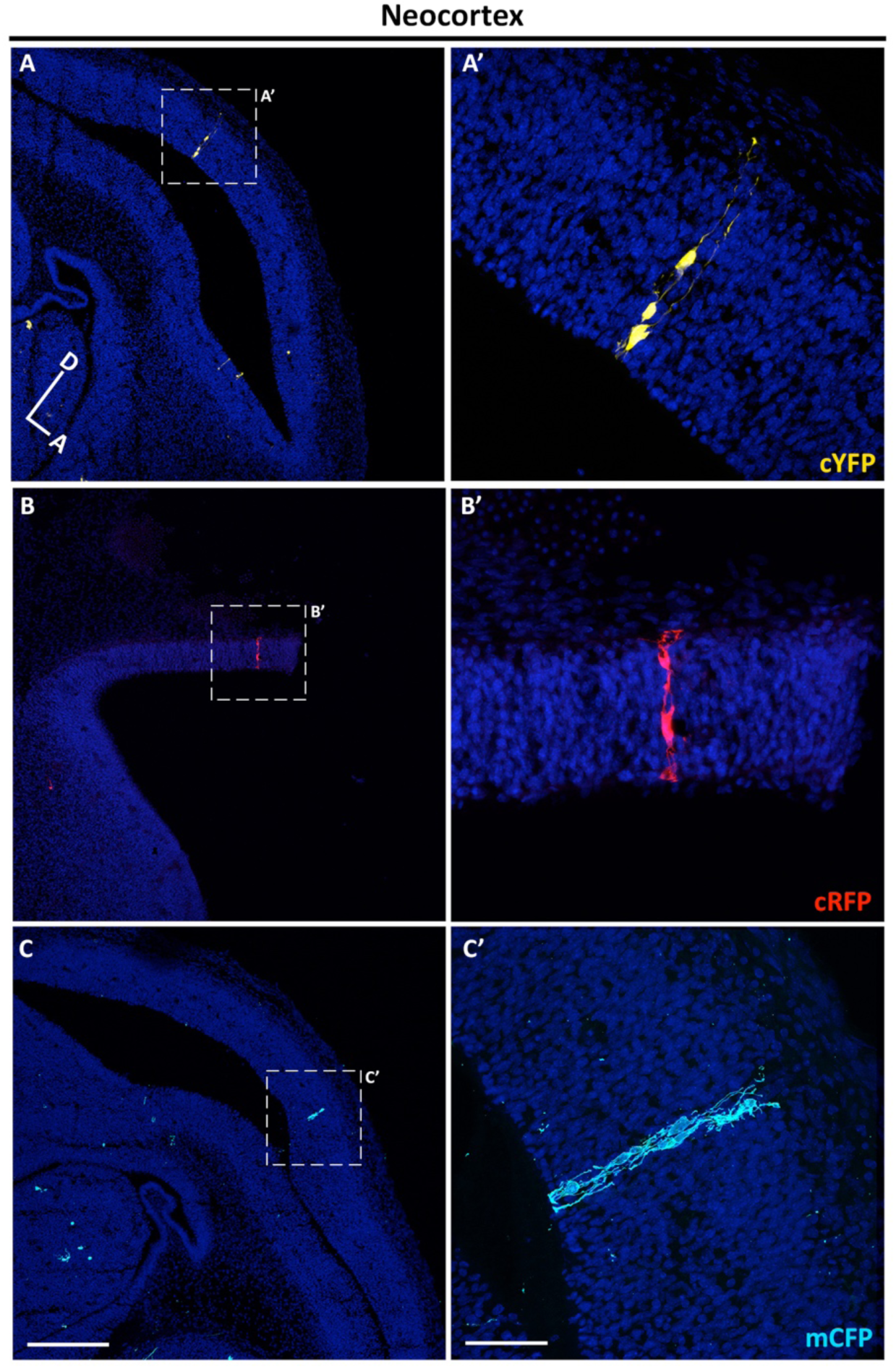
Progenitor cells labeled in neocortex with three different fluorescent proteins. (**A-C**) Confocal images of single clones labeled in the neocortex with cYFP (**A**), cRFP (**B**) and mCFP (**C**) in brain sagittal sections of E12.5 mice treated with TMX at E10.5. (**A’-C’**) High magnification images of the clones showed in **A-C**. DAPI staining (blue) reveals cell nuclei. Scale bar in C is 200 µm applies to A-C. Scale bar in C’ is 50 µm applies to A’-C’. Orientation of brains: D, dorsal; A, anterior.

**Figure 3-figure supplement 1.**
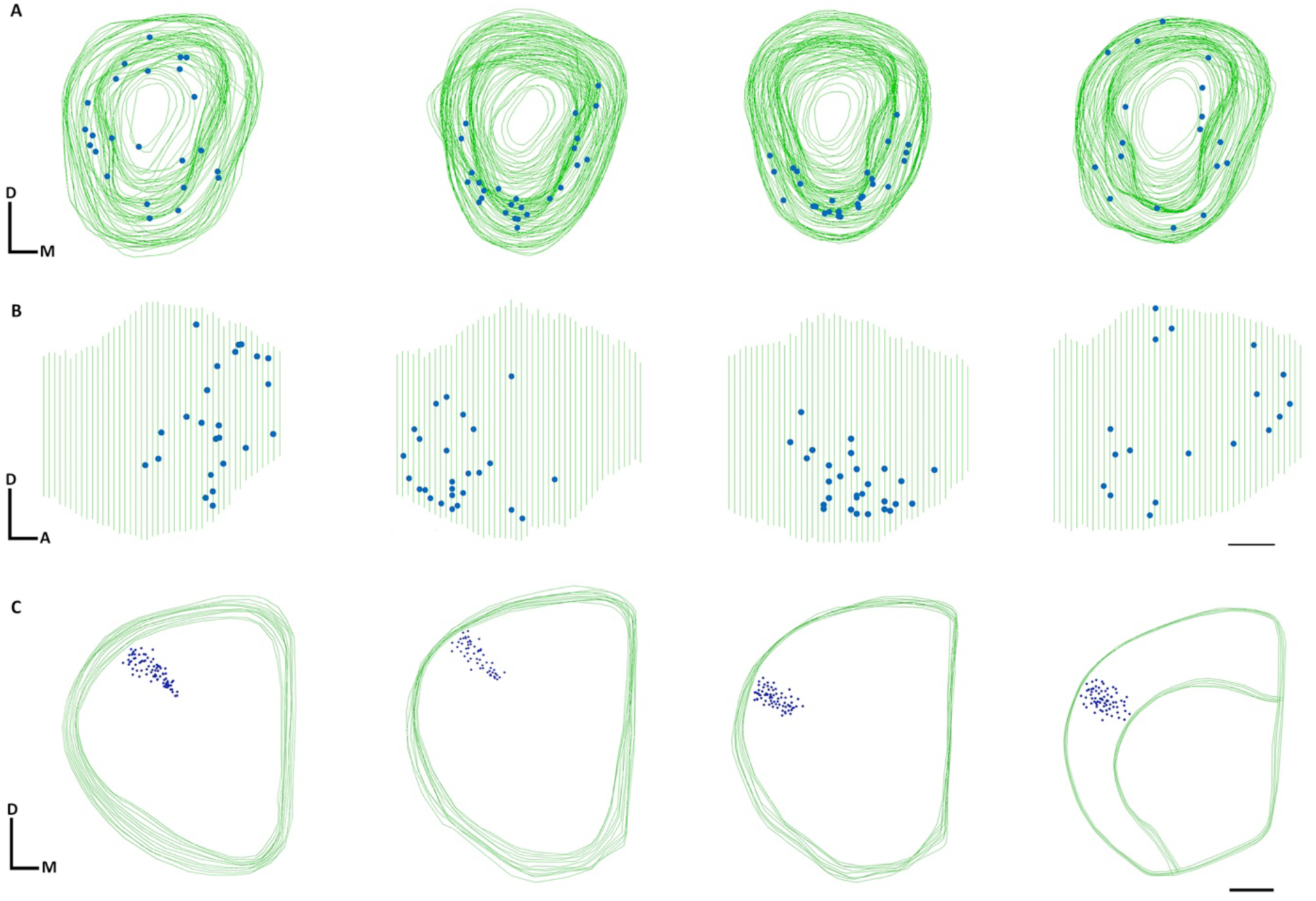
3D reconstruction of clones labelled in the olfactory bulb and neocortex. (**A-B**) 3D reconstructions from four individual clones in the OB. (**A**) Frontal and (**B**) lateral views of the OBs. (**C**) 3D reconstructions from four single clones in the neocortex. Scale bar in B is 0.5 mm and applies to A-B. Scale bar in C is 1 mm. Orientation of diagrams: D, dorsal; A, anterior; M, medial.

**Figure 3-figure supplement 2.**
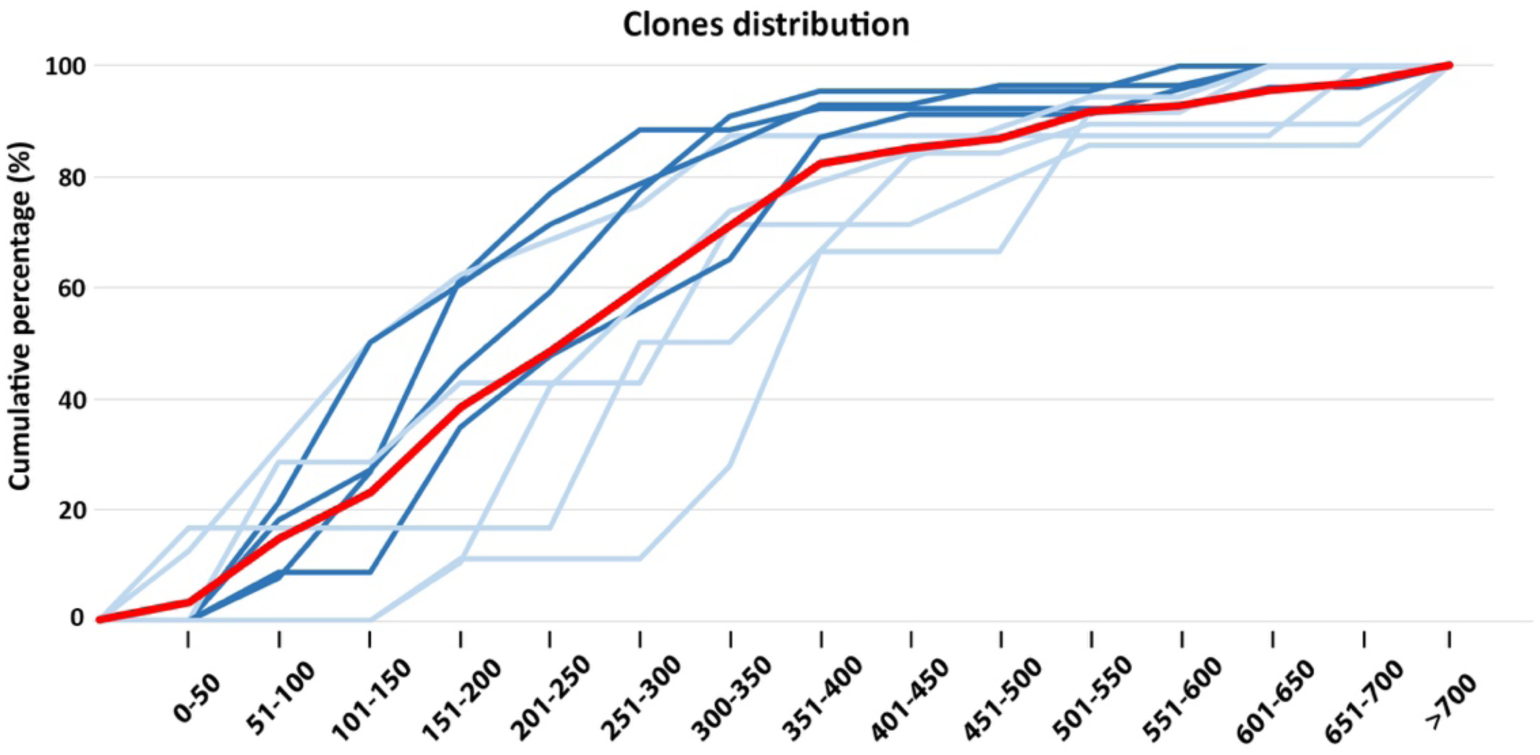
NND distribution of single clones based on their cell number. (**A**) NND cumulative percentage of individual clones analyzed in the OBs (n=9). Red line represents the NND average of all clones analyzed, while dark and light blue represent the NND of single clones containing clone sizes above or below the mean (mean=19.7), respectively.

